# Interplay of IGF1R and estrogen signaling regulates hematopoietic stem and progenitor cells

**DOI:** 10.1101/2024.03.20.585808

**Authors:** Ying Xie, Dongxi Xiang, Xin Hu, Hubert Pakula, Eun-Sil Park, Jiadong Chi, Douglas E. Linn, Luwei Tao, Zhe Li

## Abstract

Tissue stem cells often exhibit developmental stage-specific and sexually dimorphic properties, but the underlying mechanism remains largely elusive. By characterizing IGF1R signaling in hematopoietic cells, here we report that its disruption exerts sex-specific effects in adult hematopoietic stem and progenitor cells (HSPCs). Loss of IGF1R decreases the HSPC population in females but not in males, in part due to a reduction in HSPC proliferation induced by estrogen. In addition, the adult female microenvironment enhances engraftment of wild-type but not *Igf1r*-null HSPCs. In contrast, during gestation, when both female and male fetuses are exposed to placental estrogens, loss of IGF1R reduces the numbers of their fetal liver HSPCs regardless of sex. Collectively, these data support the interplay of IGF1R and estrogen pathways in HSPCs and suggest that the proliferation-promoting effect of estrogen on HSPCs is in part mediated via IGF1R signaling.

## Introduction

Stem cells in sexually dimorphic tissues (e.g., mammary gland, brain, muscle) have different properties based on their sexes^1^. However, in tissues without obvious sex differences (e.g., intestine, hematopoietic system), it has become increasingly recognized that their stem cells also exhibit sexually dimorphic functions^2,3^. The hematopoietic system is sustained by long-lived hematopoietic stem cells (HSCs), which give rise to all cells of the blood system lifelong^4^. In adults, HSCs are largely quiescent cells residing in the bone marrow (BM)^5^. Upon stimulation, HSCs go into cell cycle, produce multipotent progenitors (MPPs) and lineage-restricted hematopoietic progenitor cells (HPCs), and eventually mature blood cells. In the fetus, HSCs are proliferating cells^6,7^. The balance between HSC quiescence and proliferation are regulated by intrinsic and extrinsic factors^8^. Among extrinsic factors, estrogen increases self-renewal and proliferation of HSCs in females, by signaling through estrogen receptor alpha (ERα)^3^. However, the mechanism by which estrogen regulates HSCs remains largely elusive. A better understanding of this will have important implications for regenerative medicine and disease treatment (e.g., leukemia).

Insulin-like growth factors (IGFs, including IGF1 and IGF2) are critical regulators of cell growth, proliferation and apoptosis. Their activities are principally mediated through the insulin-like growth factor 1 receptor (IGF1R), a cell-surface tyrosine kinase and signaling molecule, to activate downstream pathways (e.g., PI3K-AKT and MAPK pathways)^9^. Both IGFs and IGF1R are commonly involved in human cancers, including leukemia^10^ and many pediatric malignancies (*e.g.*, osteosarcoma, Ewing sarcoma, and rhabdomyosarcoma)^11^. Little is known about the *in vivo* role of IGF/IGF1R signaling in hematopoiesis. Early studies suggested that IGFs might play autocrine and paracrine roles in the proliferation, survival, and differentiation of hematopoietic progenitors, particularly erythroid progenitors^12^. During development, IGF1R is expressed in fetal HSCs and the fetal liver (FL) niche may support fetal hematopoiesis by secretion of IGF2 from hepatic stromal cells^13,14^; related to this, IGF2 can expand both fetal and adult HSCs *ex vivo*^13^. In the BM niche, parathyroid hormone signaling causes a local production and release of IGF (mainly IGF1) from osteoblasts that can elicit autocrine and paracrine effects, and the primary role of IGF1 appears to promote survival of hematopoietic progenitors^15^. Conditional knockout study of IGF1R suggested that it might play a role in transition of HSCs to short-term HSCs and progenitors^16^. We showed previously that IGF/IGF1R signaling uniquely interacts with GATA1 in fetal but not adult megakaryocyte progenitors, and this developmental stage-dependent interplay may play a key role during development of megakaryocytic leukemia in Down syndrome children, when deregulated^17^.

In this study, we characterized roles of IGF1R signaling in adult and fetal hematopoiesis *in vivo*. We report a female-specific role of this pathway in regulating hematopoietic stem and progenitor cells (HSPCs) in adult mice and identify the interplay of IGF1R and estrogen signaling that modulates the proliferation of HSPCs.

## Results

### IGF1R signaling is largely dispensable for adult hematopoiesis

To study IGF/IGF1R signaling in adult hematopoiesis, we utilized an *Igf1r* conditional knockout model (*Igf1r^L^* reference ^18^) and disrupted the *Igf1r* gene by utilizing the *Vav-Cre* transgenic mouse, which leads to Cre expression specifically in hematopoietic cells^19^. Western blot confirmed that it could efficiently disrupt expression of the IGF1R protein in BM cells of *Vav-Cre;Igf1r^L/L^*mice (Fig. 1a). However, despite loss of IGF1R expression in BM hematopoietic cells, adult *Vav-Cre;Igf1r^L/L^* mice (∼8-13 weeks of age) exhibited largely normal hematopoiesis. In peripheral blood, they had normal red blood cell counts and platelet numbers, and slightly reduced (not reached statistical significance) white blood cell counts compared to those of the *Vav-Cre;Igf1r^+/+^* wild-type (WT) control mice (Supplementary Table 1). The total BM cellularity was unchanged upon IGF1R-loss (Fig. 1b). By fluorescence-activated cell sorter (FACS) analysis, we found that *Vav-Cre;Igf1r^L/L^* mice had similar percentages of myeloid, erythroid, and lymphoid populations in both their BM and spleens to those of control mice (Fig. 1c and Supplementary Fig. 1a). Furthermore, no significant difference in the frequency of Lin^-^Sca1^+^ckit^+^ (LSK) HSPC population was observed between *Vav-Cre;Igf1r^L/L^* and WT mice (Fig. 1d). Colony-forming assay showed that *Vav-Cre;Igf1r^L/L^* and WT BM cells exhibited a similar colony-forming ability (Supplementary Fig. 1b). Together, these data suggest that IGF/IGF1R signaling is largely dispensable for adult BM hematopoiesis (at least at the steady state). This observation raises a possibility that signaling through other receptor tyrosine kinases, which leads to activation of similar downstream effector pathways (e.g., PI3K/AKT, MAPK), may in part compensate for disruption of IGF1R signaling.

**Figure 1.**
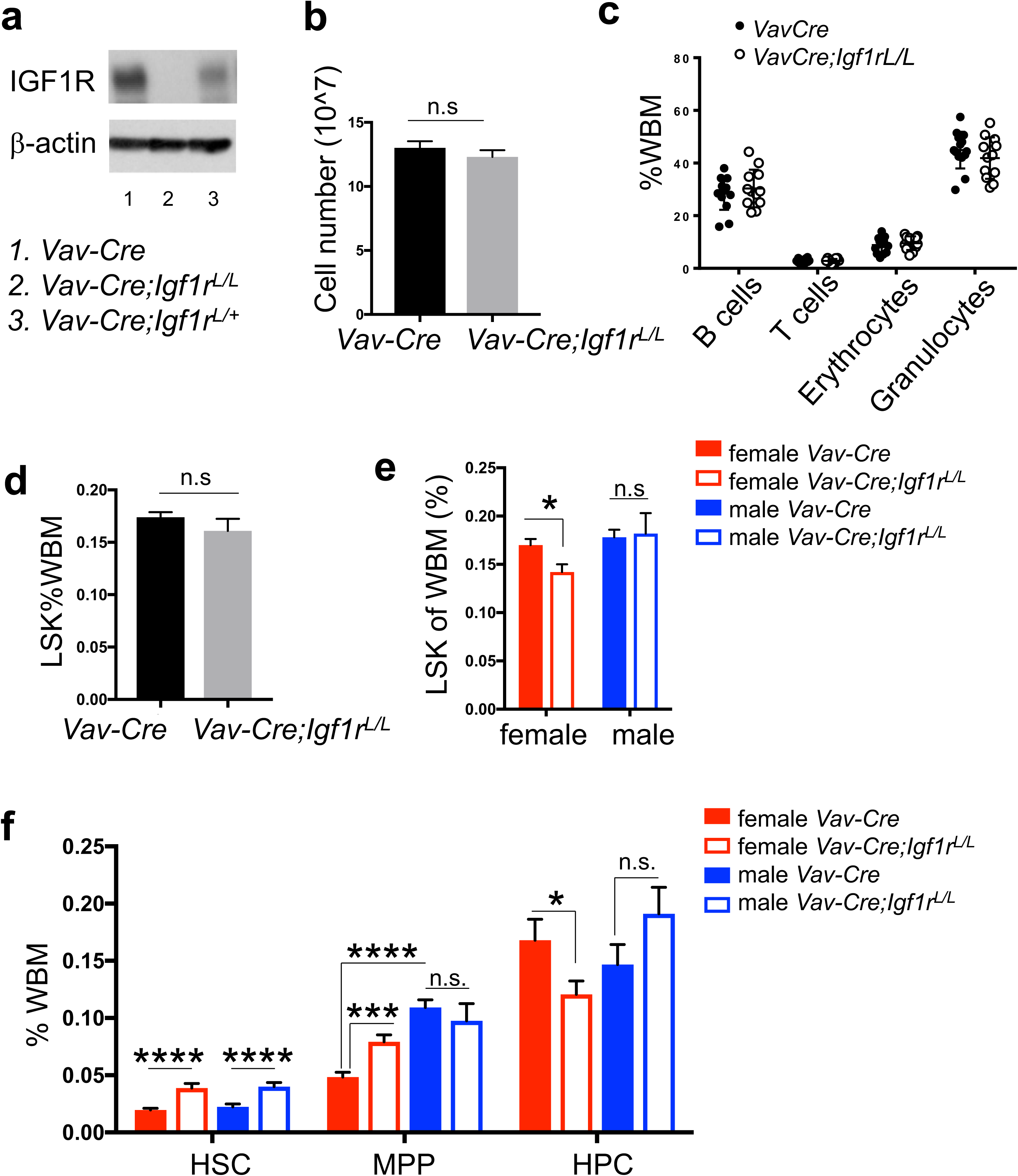
Female and male HSPCs respond differently to IGF1R-loss. (a) Western blot showing loss of IGF1R protein in BM cells from the *Vav-Cre* model. (b) Cell number of the BM cells from two tibias and two femurs per mouse. n≥11 for both WT (*Vav-Cre*) and mutant (*Vav-Cre;Igf1r^L/L^*) mice. (c) Percentages of different lineages of mature hematopoietic cells in the BM [based on whole BM (WBM)] were measured by flow cytometry (B220^+^ B cells, CD3e^+^ T cells, Ter119^+^ erythrocytes, Gr1^+^/Mac1^+^ granulocytes) between WT (*Vav-Cre*, n=14) and mutant (*Vav-Cre;Igf1r^L/L^,* n=12). (d) The LSK populations in WBM exhibited no significant difference between WT (*Vav-Cre*, n=19) and mutant (*Vav-Cre;Igf1r^L/L^,* n=19) adult mice. (e) LSK population in the BM of female or male *Vav-Cre;Igf1r^L/L^* mice compared to *Vav-Cre* control mice. n=10 for both female *Vav-Cre;Igf1r^L/L^* and *Vav-Cre* mice, n=9 for both male *Vav-Cre;Igf1r^L/L^* and *Vav-Cre* mice. (f) Percentage of each LSK HSPC subpopulation within WBM was measured by flow cytometry. n≥7 for both male and female *Vav-Cre;Igf1r^L/L^* and *Vav-Cre* mice. Data represent mean ± SEM, *p<0.05, **p<0.01, ***p<0.005, ****p<0.001, n.s.=not significant.

### Female and male HSPCs respond differently to IGF1R-loss

Thus, when grouped male and female mice together, no significant difference was observed in adult hematopoiesis between *Vav-Cre;Igf1r^L/L^* and WT mice (Fig. 1b-d and Supplementary Fig. 1a-b). Intriguingly, when we separated data from female and male mice, we observed a significant reduction in the percentages of LSK cells (i.e., HSPCs) in *Vav-Cre;Igf1r^L/L^*female mice compared to WT females (Fig. 1e). To further investigate how loss of IGF1R affects the HSPC subsets within the LSK compartment, particularly in females, we separated the BM LSK cells (HSPCs) by using SLAM markers into the CD150^+^CD48^-^LSK HSCs, CD150^-^CD48^-^LSK MPPs, and CD48^+^LSK heterogeneous restricted HPCs^20^ (Supplementary Fig. 1c). We found that both female and male *Vav-Cre;Igf1r^L/L^*mice had significantly higher percentages of HSCs compared to those of their corresponding WT control mice (Fig. 1f), consistent with a previous study^16^. In contrast, only female but not male *Vav-Cre;Igf1r^L/L^* mice had significantly lower percentages of the HPC subpopulation than those of WT controls (Fig. 1f). As HPC is the largest subpopulation within the LSK compartment (Supplementary Fig. 1c), the decrease in the percentages of LSK populations in *Vav-Cre;Igf1r^L/L^*females might be largely due to a reduction in this LSK subpopulation. In addition, we observed a significant increase in the percentage of the MPP subpopulation in *Vav-Cre;Igf1r^L/L^* females, but not in males, in relation to their WT controls (Fig. 1f).

To confirm the finding from the *Vav-Cre* model, we utilized another commonly used Cre line, *Mx1-Cre* ^21^, to disrupt *Igf1r* in BM hematopoietic cells. *Mx1-Cre* is inducible by PolyI:PolyC administration. In *Mx1-Cre;Igf1r^L/L^ ^or^ ^L/+^* mice, 3-4 weeks after the last dose of PolyI:PolyC, PCR analysis of the *Igf1r* locus confirmed extensive excision of the floxed *Igf1r* exon3 (Supplementary Fig. 1d). Upon PolyI:PolyC induction, similarly, we found there was no significant difference in the percentage of BM LSK cells in *Mx1-Cre;Igf1r^L/L^*mice and WT control mice, when female and male mice were grouped together (Supplementary Fig. 1e). However, upon separation of the data from females and males, we found this percentage was also significantly reduced in *Mx1-Cre;Igf1r^L/L^*females compared to WT control females (Supplementary Fig. 1f). Of note, in male mice, percentages of LSK cells in the *Mx1-Cre;Igf1r^L/L^* mice were increased in relation to WT control mice (Supplementary Fig. 1f). This increase might be a short-term phenotype due to PolyI:PolyC induction, as such increase was not observed in *Vav-Cre;Igf1r^L/L^* males in which deletion of *Igf1r* has occurred in hematopoietic cells since the fetal stage^19^ (Fig. 1e).

### IGF1R-loss abolishes estrogen-triggered proliferation increase in HSPCs

We next investigated whether the female-specific decrease of HSPCs in *Vav-Cre;Igf1r^L/L^* mice was due to any change in cell survival or proliferation, by measuring their AnnexinV^+^ or Ki67^+^ fractions, respectively (Supplementary Fig. 2a). Compared to males, HSPCs in females exhibited increased proliferation (Fig. 2a), consistent with the previous finding^3^. Intriguingly, loss of IGF1R in HSPCs in females led to a significant reduction in their proliferative fraction, to a level comparable to that of WT male HSPCs (Fig. 2a). In contrast, loss of IGF1R in males did not lead to any significant change in the percentage of Ki67^+^ cells in their LSK HSPC compartment (Fig. 2a). Furthermore, loss of IGF1R in both female and male hematopoietic cells led to increases in their AnnexinV^+^ apoptotic cells, in both the LSK and the entire BM compartments (Supplementary Fig. 2b).

**Figure 2.**
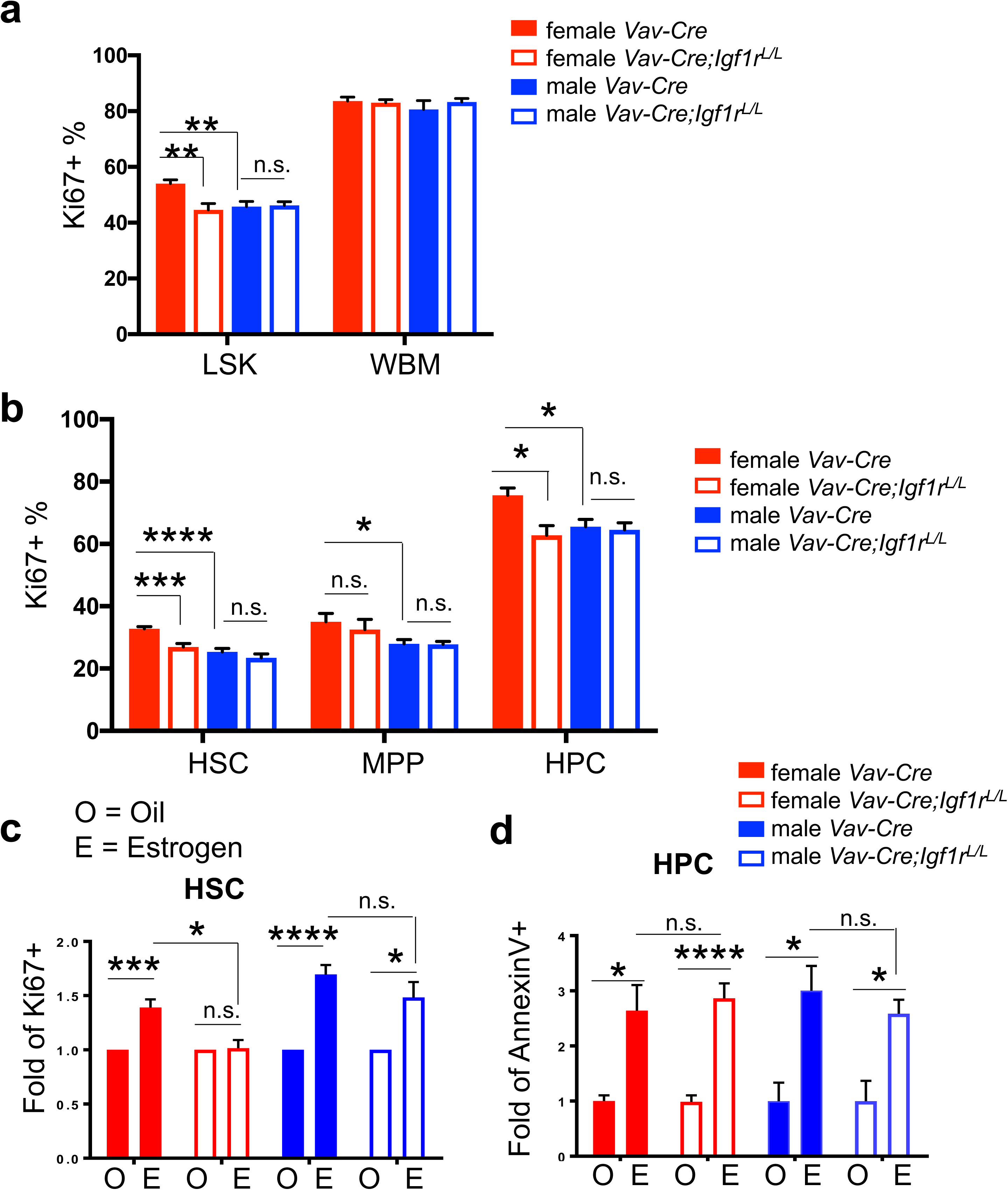
Increased HSPC proliferation triggered by estrogen in female mice is abolished upon IGF1R-loss. (a) Percentages of Ki67^+^ cells were measured for both LSK and WBM populations to reflect cells in cell cycle. n≥5 for both male and female *Vav-Cre;Igf1r^L/L^* and *Vav-Cre* mice. (b) Percentages of Ki67^+^ proliferating cells were measured for all LSK subpopulations. n≥5 for both male and female *Vav-Cre;Igf1r^L/L^* and *Vav-Cre* mice. (c) Percentages of Ki67^+^ proliferating cells were measured for HSCs of female or male mice with the indicated genotypes (legend the same as in d) treated with oil (O) or β-estradiol (E), and are shown as fold changes in relation to those of corresponding controls (i.e., oil-treatment). n≥3 for both male and female *Vav-Cre;Igf1r^L/L^* and *Vav-Cre* mice. (d) Percentages of AnnexinV^+^ apoptotic cells were measured for HPCs of female or male mice with the indicated genotypes treated with oil (O) or β-estradiol (E), and are shown as fold changes in relation to those of corresponding controls (i.e., oil-treatment). n≥3 for both male and female *Vav-Cre;Igf1r^L/L^* and *Vav-Cre* mice. Data represent mean ± SEM, *p<0.05, **p<0.01, ***p<0.005, ****p<0.001, n.s.=not significant.

Next, we analyzed the proliferative and apoptotic fractions in HSPC subpopulations. In all three HSPC populations (HSC, MPP, HPC) of WT mice, females had significantly higher proliferative fractions than those of males (Fig. 2b). In contrast, only the HPC subpopulation in WT females had significantly more apoptotic cells than males (Supplementary Fig. 2c). Upon loss of IGF1R, in females we observed significant reduction of Ki67^+^ proliferative cells and significant increase of AnnexinV^+^ apoptotic cells in both HSC and HPC subpopulations (Fig. 2b and Supplementary Fig. 2c); in contrast, in males we only observed significant increase of AnnexinV^+^ apoptotic cells in all their HSPC subpopulations (Supplementary Fig. 2c), and there was no change in the proliferation of HSPC subsets (Fig. 2b).

To further investigate the role of IGF1R signaling in estrogen-mediated HSC proliferation and apoptosis, we administered β-estradiol (E2) to young adult female and male mice for 1 week. The level of estrogen in female mice upon injection is comparable to that of females during pregnancy, while the level in the injected male mice is comparable to that of adult females ^3^. As expected, E2 treatment increased the numbers of proliferating HSCs in both female and male WT mice (Fig. 2c and Supplementary Fig. 2d). Loss of IGF1R largely abolished this E2-induced increase in the proliferation of female HSCs, and also slightly reduced this proliferation increase in male HSCs (Fig. 2c and Supplementary Fig. 2d). E2 treatment did not significantly change proliferation of other LSK subpopulations (i.e., MPP, HPC) in both female and male mice (except downregulation of proliferation of the MPP and HPC subsets in females with IGF1R-loss, Supplementary Fig. 2d). In contrast, E2 treatment significantly increased apoptosis of HPCs in both sexes, regardless of whether *Igf1r* is WT or null (Fig. 2d and Supplementary Fig. 2e). E2 treatment did not significantly change apoptosis of the HSC and MPP subsets in both *Igf1r-WT* and *Igf1r-null* female or male mice (except increased apoptosis of HSCs and MPPs in WT males, Supplementary Fig. 2e).

### Molecular mechanism for the interplay of IGF1R and estrogen signaling

So far, our data suggest that in adults, IGF1R signaling plays a key role in mediating estrogen-induced increase in the proliferation of HSCs, and probably also HPCs; IGF1R signaling also plays a role in the survival of HSPCs, in particular, HPCs, regardless of the estrogen status (Fig. 2). To gain insights into the molecular mechanism underlying this complex phenotype, we retrospectively analyzed publicly available expression profiling data for IGF1R and estrogen signaling in HSCs^3,16^. We generated gene sets for genes upregulated or downregulated upon E2 treatment in female HSCs compared to male HSCs^3^, as well as those for genes upregulated or downregulated in *Igf1r-null* HSCs compared to WT HSCs^16^. We found that genes involved in the PI3K-AKT pathway are either only significantly upregulated in female HSCs or significantly upregulated in both male and female HSCs (upon E2 treatment) in relation to mock controls (Supplementary Fig. 3a-b). This analysis suggests that PI3K-AKT may be a key pathway to mediate the pro-proliferation effect of estrogen signaling on HSCs, especially female HSCs, possibly via ERα^3^ (Fig. 3a). Of note, the PI3K-AKT pathway is also a key downstream effector pathway to mediate the effect of IGF1R signaling on cellular proliferation and survival^9^. IGF1R signaling can positively regulate estrogen signaling output via the PI3K-AKT pathway; this can be achieved by increasing ERα activity via phosphorylation and/or by enhancing translation of ERα-mediated genes through increasing mRNA cap dependent translation^22^. We thus propose that estrogen signaling enhances proliferation of HSCs through the IGF1R signaling, possibly via its downstream PI3K-AKT effector pathway and by modulating the transcriptional and/or translational output of ERα (Fig. 3a).

**Figure 3.**
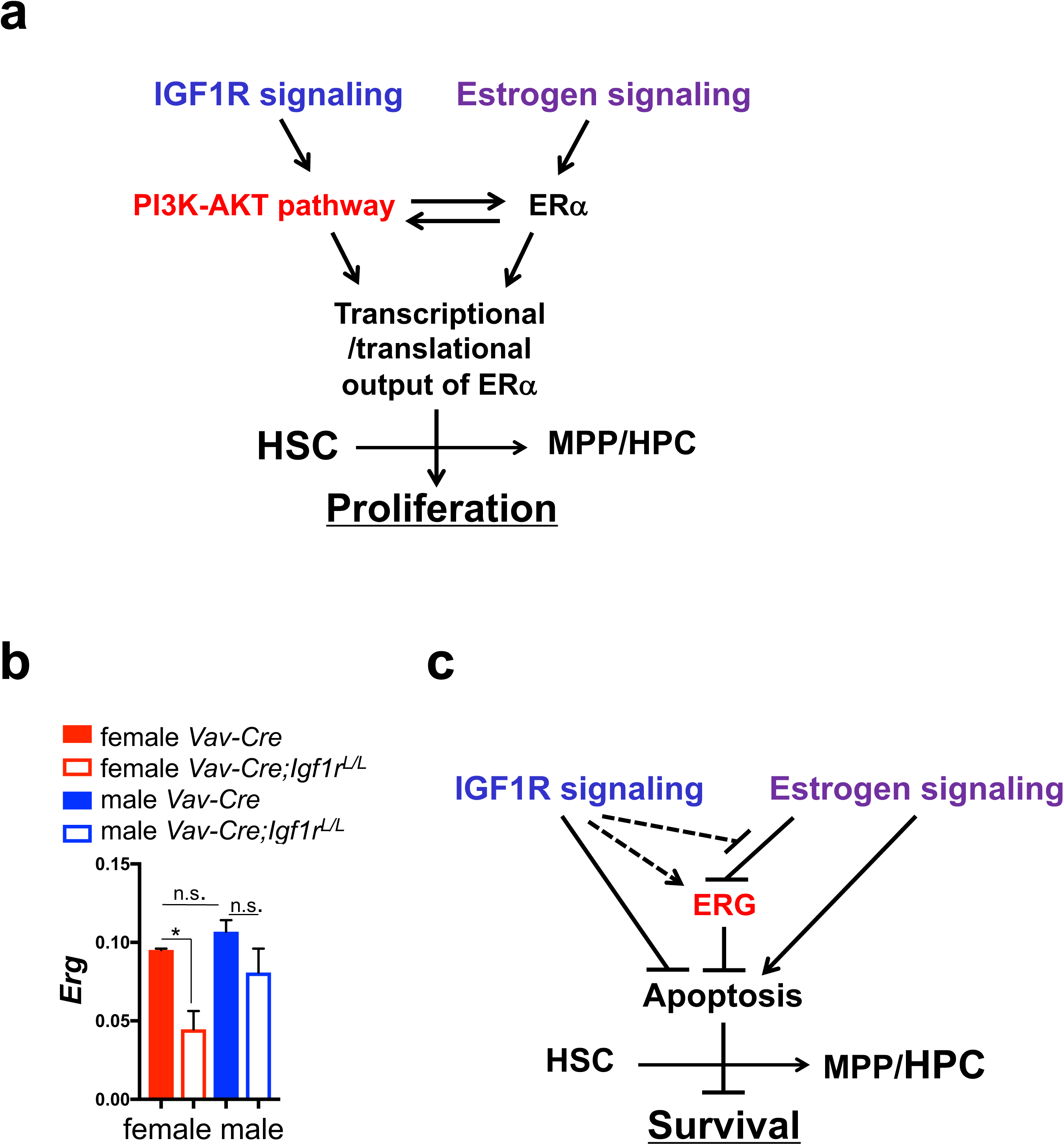
Models for the molecular mechanisms underlying the interplay of IGF1R and estrogen signaling. (a) Schematic diagram showing estrogen signaling and IGF1R signaling regulate proliferation of HSPCs via the PI3K-AKT pathway. (b) qRT-PCR analysis showing relative expression levels of *Erg* in female and male BM HSCs with the indicated genotypes. n≥3. Data represent mean ± SEM, *p<0.05, n.s.=not significant. (c) Schematic diagram showing estrogen and IGF1R signaling pathways exhibit different effects on apoptosis of HSPCs in part via their opposite regulations of ERG.

We measured expression levels of several HSC-related transcription factor (TF) genes. We found that whereas the level of *Esr1* itself was unchanged in female and male HSCs upon loss of IGF1R (Supplementary Fig. 3c), the level of a TF gene, *Erg*, was significantly reduced in *Vav-Cre;Igf1r^L/L^*females in relation to WT control females; its level was also slightly reduced in *Vav-Cre;Igf1r^L/L^* males compared to WT males (Fig. 3b). ERG is a key TF for maintenance and survival of HSCs^23,24^. Of note, the HSPC phenotype we observed in *Vav-Cre;Igf1r^L/L^* females is similar to that of an *Erg* knockdown mouse model we reported previously^24^, raising a possibility that the *Igf1r-null* HSPC phenotype may be in part mediated by downregulation of *Erg*. In particular, we showed that ∼40% reduction in *Erg* dosage led to a reduction in the number of LSK HSPCs, in part due to their increased apoptosis^24^. Interestingly, it was reported previously that the transcriptional activity of ERG can be repressed by ERα^25^. We and others have shown that estrogen signaling can increase apoptosis of HSPCs (Fig. 2d and Supplementary Fig. 2b-c,e, and references ^26,27^). We thus propose that estrogen signaling may increase apoptosis of HSPCs in part via repression of ERG, whereas IGF1R signaling may suppress HSPC apoptosis in part via upregulation of the transcriptional activity of ERG; the latter may be achieved either directly (e.g., via upregulation of *Erg* expression) or indirectly via suppression of the repressive effect of estrogen signaling on ERG (Fig. 3c). The indirect model may explain why in female HSCs, loss of IGF1R exhibited a more profound effect on downregulation of *Erg* expression than in males (Fig. 3b).

### Female microenvironment increases contribution of WT but not *Igf1r-null* HSCs

So far, we have demonstrated the role of IGF1R signaling in mediating estrogen-induced proliferation of HSCs, as well as a largely gender/estrogen-independent role of this pathway in the survival of HSPCs. HSCs reside in the BM and its function is affected by the microenvironment. Estrogen and IGFs are both environmental factors affecting hematopoiesis. To further understand their interplay for regulating HSCs function, we performed a competitive BM transplantation assay. Irradiated *Rag2^-/-^* female or male recipient mice were transplanted with the same numbers of CD45.1 competitor BM cells together with CD45.2 donor BM cells from either *Mx1-Cre;Igf1r^L/L^;Rosa26^LSL-YFP^* (*R26Y*) or *Mx1-Cre;R26Y* (WT) female mice. In these mice, Cre-mediated recombination disrupts *Igf1r* and meanwhile turns on the YFP reporter. Five days after transplantation, we measured the initial contribution of the CD45.2 cells as a starting point. We then treated the mice with PolyI:PolyC to induce Cre-mediated disruption of *Igf1r*. After the last dose of PolyI:PolyC, we measured the percentage of YFP^+^ cells as the contribution of donor cells to hematopoiesis in recipients at week 4, 8, 12, and 16 (Fig. 4a). We used the inducible *Mx1-Cre* model to avoid any potential homing difference between *Igf1r-null* and *Igf1r-WT* cells.

**Figure 4.**
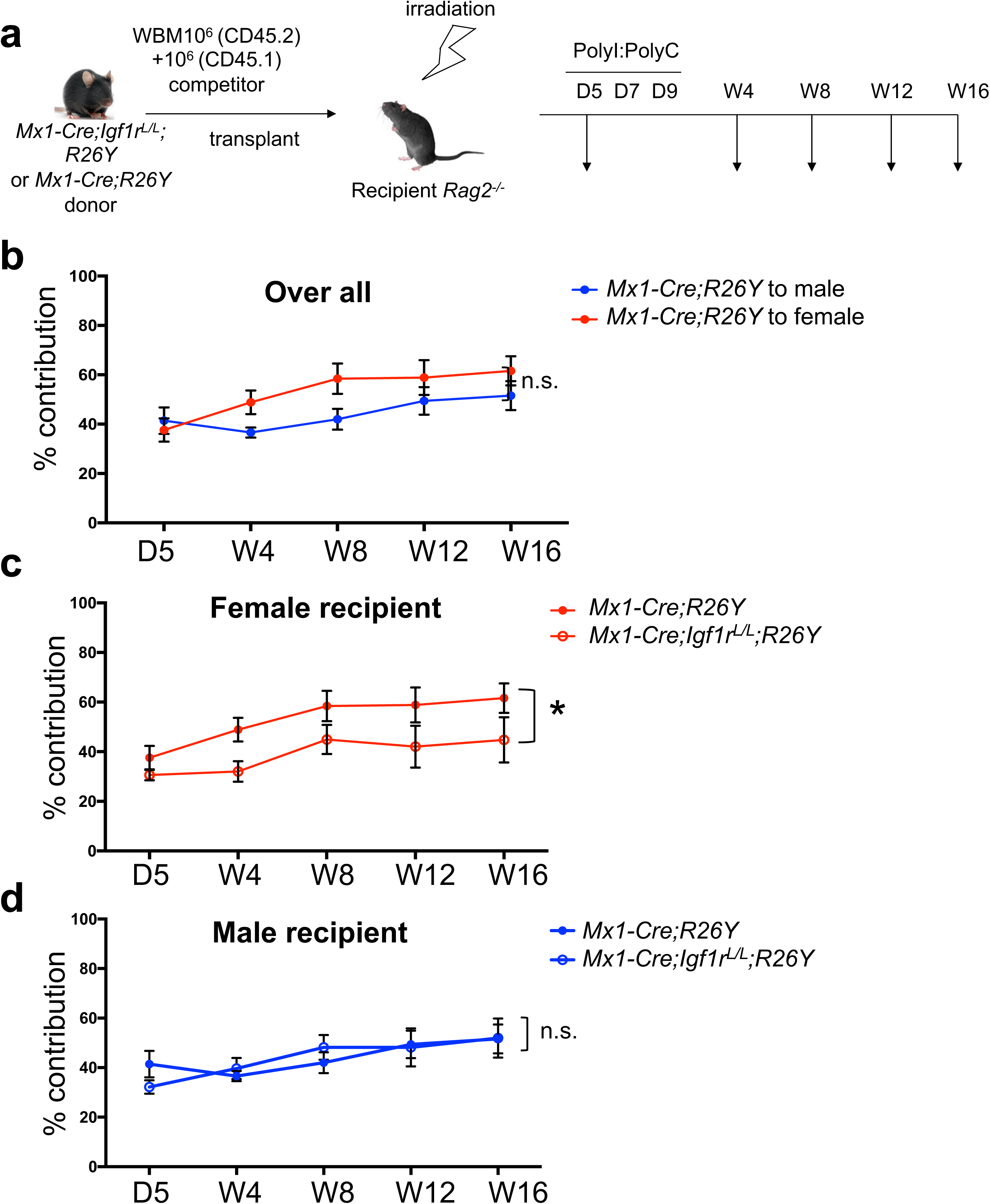
Female microenvironment increases contribution of WT but not *Igf1r*-null HSCs. (a) Schematic diagram showing experimental design of the competitive BM transplantation. D: day; W: week. (b) Donor contribution (based on percentage of CD45.2^+^YFP^+^ cells) from *Mx1-Cre;R26Y* (WT) cells in peripheral blood was measured over time in female versus male recipient mice. (c-d) Donor contribution based on percentage of CD45.2^+^YFP^+^ cells over time in peripheral blood was measured in both female (c) and male (d) recipient mice. n≥3 for each data point. Data represent mean ± SEM. Pearson’s correlation coefficient analysis was used to calculate the *P* values for the whole trend. *p<0.05, n.s.=not significant.

Although not reached statistical significance, we observed that the same donor cells exhibited better engraftment in female recipients than male recipients (Fig. 4b). Intriguingly, upon induced IGF1R-loss, the *Igf1r-null* donor cells exhibited a significant reduction in their engraftment in female recipients, but not in male recipients, when compared to *Igf1r-WT* donor cells (Fig. 4c-d). Of note, IGF1R-loss in donor HSCs did not abolish their ability to repopulate the hematopoietic system in female recipients, but rather reduced their engraftment to a level comparable to that in male recipients (Fig. 4c-d). A poorly understood observation in HSC transplantation is the enhanced engraftment of human HSCs in female compared to male mice^28^. Estrogen has been proposed as a factor contributing to this^1^. Although irradiation is expected to reduce estrogen expression in female mice by damaging ovarian function (at least at the initial stage post-irradiation), it was reported that total body γ irradiation actually led to a significant increase in the serum estradiol level two months after irradiation and the increase was sustained to at least 12 months post-irradiation^29^. As the majority of our analysis time points were at two months or more after irradiation, our engraftment data in female versus male recipients is consistent with a model in which estrogen can enhance HSC engraftment and IGF1R signaling is a key player in mediating this.

We also analyzed the contribution of *Igf1r-null* and *Igf1r-WT* donor cells to different hematopoietic mature lineages in female and male recipients. The contribution to myeloid cells (Gr1^+^/Mac1^+^) was slightly reduced in both female and male recipients upon IGF1R-loss (Supplementary Fig. 4a), whereas the contribution to T cells (CD3e^+^) exhibited no significant difference (Supplementary Fig. 4b). Of note, the B lymphocyte-lineage (B220^+^ cells) had the most notable changes upon IGF1R-loss. Loss of IGF1R significantly reduced the B cell-lineage reconstitution in both female and male recipients, but with a more dramatic reduction in female recipients (Supplementary Fig. 4c-d). This phenotype may be related to an additional role of IGF1R signaling in stimulating differentiation and proliferation of B cells^30^. In addition, although it was reported that estrogen negatively regulates B lymphopoiesis^31,32^, due to the higher overall hematopoietic contribution of HSPCs in female than male hosts (Fig. 4b), the difference of B lineage contribution in female versus male hosts would be minimized. However, when *Igf1r* was disrupted, the gender-specific reduction in overall hematopoietic contribution in female hosts would make the reduction in the B lineage appear greater in female than male recipients.

### Loss of IGF1R affects both female and male fetal hematopoiesis

We demonstrated that IGF1R signaling plays a key role in mediating estrogen-induced increase in proliferation of HSPCs in adult females. Another important estrogen-rich environment for HSPCs can be found during fetal development. At this stage of development, HSPCs in both female and male fetuses are equally exposed to estrogens in their circulation, which are released from the placental trophoblasts^33,34^. Although it remains unanswered as to whether estrogens play a role in the rapid cycling and self-renewal proliferation of fetal HSPCs, the interplay of estrogen and IGF1R signaling in adult HSPCs raises a possibility that loss of IGF1R may have a more notable impact on fetal HSPCs in both female and male fetuses. To test this, we used *Tie2-Cre* to disrupt *Igf1r* in hematopoietic (and endothelial) cells in fetuses^24^, as disruption of *Runx1* (required for the birth of HSCs) by this Cre line led to a complete loss of definitive hematopoiesis^35^. Western blot analysis confirmed a complete loss of IGF1R protein in fetal hematopoietic cells from *Tie2-Cre;Igf1r^L/L^* fetuses (Supplementary Fig. 5a). By timed breeding, we analyzed FL HSPCs from both female and male E13.5 fetuses. By FACS analysis (Supplementary Fig. 5b), we found that the percentages of Lin^-^Sca1^+^ckit^+^ (LSK) HSPCs in FLs from both female and male *Tie2-Cre;Igf1r^L/L^* mice were significantly reduced compared to their corresponding WT controls (Fig. 5a). Furthermore, by hematopoietic CFU assay, we found that the *Tie2-Cre;Igf1r^L/L^* FLs contained significantly less colony-forming cells compared to WT FLs, regardless of the sex (Fig. 5b). Related to this impaired fetal hematopoiesis upon IGF1R-loss, we observed a slight reduction in the frequency of *Tie2-Cre;Igf1r^L/L^*mice after birth (i.e., less than the expected Mendelian ratio, Supplementary Fig. 5c). Together, these data suggest that different from adult hematopoiesis, IGF1R signaling may play a key role in proliferation and/or survival of fetal HSPCs, regardless of the sex; although not definitively proven, we propose that this role of IGF1R signaling may be also interconnected with estrogen signaling.

**Figure 5.**
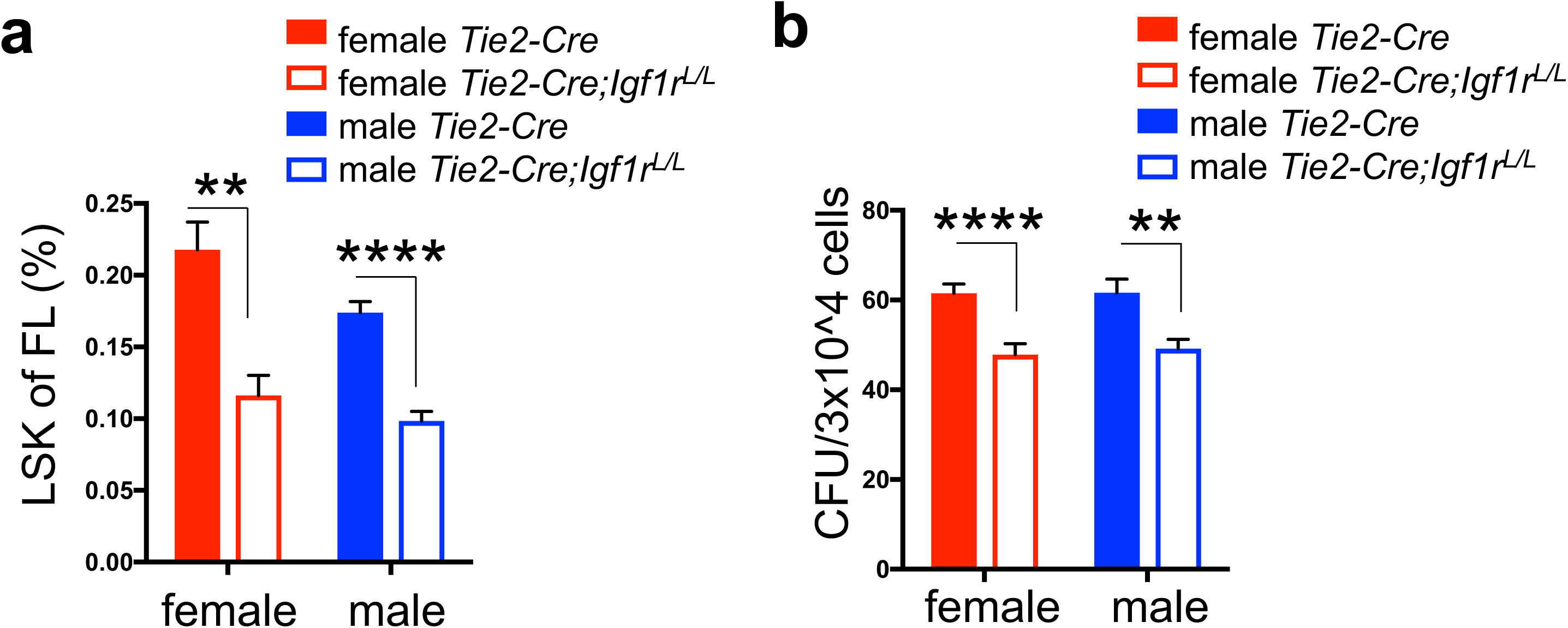
Potential interplay of IGF1R and estrogen signaling in fetal HSPCs. (a-b) LSK percentage (a) and colony forming units (CFU) (b) were measured in both male and female E13.5 FLs. n≥3 for LSK percentage and n≥6 for CFU assay. Data represent mean ± SEM. **p<0.01, ****p<0.001.

## Discussion

In this study, we revealed several phenotypes of HSPCs associated with IGF1R-loss, including an increase in the HSC subpopulation, increased apoptosis in all BM HSPC subsets, reduction in the proliferative portions of HSCs and HPCs in adult females, and reduction in FL HSPCs in both female and male fetuses. Among these, the latter two may be related to the interplay of estrogen and IGF1R signaling pathways. Our data support that: 1) under the physiological setting when there is an increased demand for the output from the hematopoietic system (e.g., in females during pregnancy, or possibly also in fetuses during development), IGF/IGF1R signaling appears to play important roles; 2) estrogen may play a key role in these settings and IGF/IGF1R signaling is particularly important for supporting this estrogen-mediated process.

Of note, in the estrogen-treatment experiment, we found that loss of IGF1R exhibited a more profound effect in “canceling out” increased proliferation of HSCs triggered by E2 in females than males (Fig. 2c and Supplementary Fig. 2d). The biological difference between female and male is not only due to an endocrine difference, but also attributed to sex-specific genetic architecture^36^; that is, sex can be viewed as a variant equivalent to environmental factors, which interacts with the genotype (e.g., loss of IGF1R), leading to sex-specific phenotypic differences, in addition to those caused by the endocrine difference (e.g., estrogen) alone. In support of this, a retrospective analysis of published gene expression data for male and female HSCs treated with or without E2 (based on reference ^3^) revealed that among all upregulated or downregulated genes in male and female HSCs upon E2 treatment, only a small number of genes were in common (Supplementary Fig. 6a). Furthermore, we compared molecular outcomes of E2 treatment on female and male HSCs (based on reference ^3^) to those from HSCs with IGF1R-loss (based on reference ^16^); by gene set enrichment analysis (GSEA)^26,37^, we found that for genes positively correlated with IGF1R signaling, E2 treatment largely upregulated their expression in both female and male HSCs, whereas for genes negatively correlated with IGF1R signaling (i.e., *Igf1r* downregulated genes), E2 treatment largely downregulated their expression in female HSCs but not in male HSCs (Supplementary Fig. 6b-c). These analyses suggest that there is a difference at the molecular level for responses of female and male HSCs to modulations of estrogen and IGF1R signaling, with female HSCs exhibiting more concordant estrogen signaling and IGF1R signaling-related gene expression changes, thus further supporting a notion that the female-specific genetic architecture may be more tightly wired with the IGF/IGF1R pathway to support female-specific physiology.

Similar to the IGF/IGF1R pathway, multiple developmental pathways (e.g., Notch, Wnt, and Hedgehog signaling) also play important roles in hematopoiesis during development, but are dispensable for adult hematopoiesis^38–43^. The underlying mechanism for this developmental stage-specific dependency remains largely elusive. Our study of IGF/IGF1R signaling and its potential link to estrogen signaling (e.g., during fetal hematopoiesis) provide one possible mechanism to explain this. Of note, although these signaling pathways are largely dispensable for adult hematopoiesis, they are often utilized by leukemic cells for their survival and proliferation, when aberrantly activated^10,44–46^. A better understanding of the molecular basis for their differential requirements at different stages of hematopoietic development may offer a novel therapeutic opportunity to treat such leukemia.

The interplay of estrogen and IGF/IGF1R signaling is also conserved in other settings, particularly in malignancies such as breast cancer^22,47^. IGF/IGF1R signaling can regulate ERα activity by altering its phosphorylation and activation^22,47^; it can also enhance estrogen signaling by increasing translation of ER-mediated genes^22^. Conversely, estrogen signaling can affect IGF/IGF1R signaling as well^22,47^. For instance, E2 can increase expression of IGF1R, decrease IGF2R expression, stimulate phosphorylation of IGF1R, and influence expression and activity of other components in the IGF/IGF1R pathway. We propose that in HSPCs, the interplay of these two pathways may be mediated in part via the PI3K-AKT pathway (e.g., to support HSPC proliferation) and the hematopoietic TF, ERG (e.g., to support survival of HSPCs) (Fig. 3). In leukemia, we reported previously that IGF/IGF1R signaling plays a key role in the development and/or maintenance of Down syndrome acute megakaryoblastic leukemia (DS-AMKL), which has a fetal origin, and may represent a potential therapy target to treat this malignancy^17^. From the current study, we observed the interplay of estrogen and IGF signaling in regulating HSPCs in adult females and possibly also in fetuses. This raises a possibility that leukemic cells that are highly dependent on IGF/IGF1R signaling may be also sensitive to inhibition of estrogen signaling, and that inhibition of both estrogen and IGF/IGF1R signaling may have a synergistic effect. As DS-AMKL and its precursor lesion DS transient leukemia (DS-TL) are pediatric leukemia/pre-leukemia with a fetal origin, we suspect placental estrogens may play a role in the pathogenesis of DS-TL and that a dramatic drop in the estrogen level right after birth may in part explain the spontaneous remission of DS-TL shortly after birth.

## Methods

### Mice

Mouse strains used in this study were obtained from the Jackson Laboratory (JAX, Bar Harbor, ME), including: *Igf1r* conditional knockout (*Igf1r^L^*, JAX Stock No: 012251), *Mx1-Cre* transgenic mouse (JAX Stock No: 003556), *Tie2-Cre* transgenic mouse (JAX Stock No: 004128), *Rosa26^LSL-YFP^* (*R26Y*) reporter mouse (JAX Stock No: 006148), B6.SJL (CD45.1) mouse (JAX Stock No: 013591); in addition, *Vav-Cre* transgenic mouse and *Rag2^-/-^* mouse are gifts from Laboratory of Dr. Stuart Orkin (BCH). All animal procedures and protocols were approved by the institutional Animal Care and Use Committee (IACUC) of Boston Children’s Hospital. *Rag2^-/-^* mice were used for competitive bone marrow transplantation assays. PolyI:polyC (invivogen, 31852-29-6) was i.p. injected to mice for inducing *Mx1-Cre*. Mice were sacrificed for analysis three to four weeks after the last injection. *Vav-Cre* mice with ages of 8-13 weeks were used for adult study. *Tie2-Cre;Igf1r^L/+^* male mice were mated with *Igf1r^L/+^* female mice by timed breeding to generate E13.5 fetuses. β-estradiol (E2, Sigma-Aldrich, E1132) was administered daily (2μg/day) through i.p. injection for seven days before analysis.

### Complete Blood Count (CBC) analysis

Peripheral blood was collected with EDTA-coated capillaries (BD Biosciences, San Jose, CA). Complete blood count was performed as described^17,48^. Blood samples (∼250μl) were obtained by retro-orbital sinus bleeding and analyzed using an automated system (Advia 120, Bayer).

### Flow Cytometric Analysis

BM, peripheral blood (PB), and FL single cell suspensions were prepared and flow cytometry was performed as described^17,48^ using an Accuri C6 (BD Biosciences) analyzer, or DXP11 analyzer (Cytek, Fremont, CA) or the FACSAria sorter (BD Biosciences). Sorting was performed using a FACSAria sorter. The following antibodies were used: CD3e-biotin, B220-biotin, Mac1-biotin, Gr1-biotin, Ter119-biotin were used as lineage for adult BM; CD3e-biotin, B220-biotin, Gr1-biotin, Ter119-biotin were used as lineage for FLs. Streptavidin-eFluo700, Sca1-APCcy7, c-kit-PerCPcy5.5, CD150-PE, and CD48-APC were used for HSPCs. CD3e-PE, B220-APC, Mac1-PEcy7, Mac1-PE, Gr1-APCcy7, Ter119-biotin, Ter119-APC, and Streptavidin-PerCP, were used to study mature lineages. CD45.1-PEcy7 and CD45.2-PerCPcy5.5 were used for competitive BM transplantation. AnnexinV-PEcy7, Ki67-PEcy7, and DAPI were used for measuring apoptosis or proliferation. All antibodies were from eBioscience or BD Biosciences.

### Colony-Forming Assays

FL and BM colony forming assays in methylcellulose (MethoCult M3434, Stemcell Technologies, Vancouver, Canada) were performed according to the manufacturer’s instructions.

### Western Blot Analysis

Cells were lysed and Western blot analysis was performed as described^49^. Western blots were probed with IGF1R antibody (Santa Cruz, CA) and anti-β-Actin antibody (Thermo Fisher Scientific, Waltham, MA) was used as a loading control.

### Apoptosis and Proliferation Analysis

BM single cell suspensions were prepared and flow cytometry was performed as described^17,48^, using a FACSAria sorter (BD Biosciences). For apoptosis analysis, cells were stained with Annexin-PEcy7 first. After washing, cells were stained with biotinylated lineage markers. After another wash, cells were stained with streptavidin-eFluo700, Sca1-APCcy7, c-kit-PerCPcy5.5, CD150-PE, and CD48-APC. After washing, cells were suspended in PBS with 4’,6-diamidino-2-phenylindole (DAPI) and subjected to flow cytometric analysis. For proliferation analysis, cells were stained with biotinylated lineage markers. After wash, cells were stained with streptavidin-eFluo700, Sca1-APCcy7, c-kit-PerCPcy5.5, CD150-PE, and CD48-APC. After washing, cells were fixed with the fixation/permeabilization kit (eBiosciences). After fixation and washing, cells were stained with Ki67-PEcy7. After washing, cells were suspended in PBS with 4’,6-diamidino-2-phenylindole (DAPI) and subjected to flow cytometric analysis.

### Quantitative RT-PCR

Total RNAs from sorted HSCs were purified by total RNA isolation with TRIzol (Thermo Fisher Scientific, Waltham, MA). cDNA was generated with iScript (Bio-Rad, Berkeley, CA) according to the manufacture’s protocol. Quantitative RT-PCR (qRT-PCR) was performed using FastStart SYBR Green Master kit (Roche, Indianapolis, IN).

### Competitive Bone Marrow Transplantation

Competitive bone marrow transplantation is described in Figure 4a. Briefly, ∼1×10^6^ BM cells were obtained from either *Mx1-Cre;Igf1r^L/L^;R26Y* or *Mx1-Cre;Igf1r^+/+^;R26Y* female mice (CD45.2), mixed with an equal amount of BM cells from B6.SJL (CD45.1) mice, and injected together into lethally irradiated *Rag2^-/-^* male or female mice (CD45.1/CD45.2). Five days after transplantation, initial contribution of the CD45.2 cells were measured as a starting point. PolyI:PolyC was i.p. injected into the recipient mice at a dose of 12.5μg/g body weight/day, once every other day, three times in total. The percentage of YFP^+^ cells was measured (as the level of contribution from donor cells) at week 4, 8, 12, and 16 after the last dose of PolyI:PolyC.

### Analysis of datasets

The microarray data for E2-treated HSCs (compared with oil-treated HSCs) were obtained from Gene Expression Omnibus under accession number GSE52711^3,16^; the RNAseq data comparing *Igf1r*-null HSCs to WT HSCs were obtained from ArrayExpress under accession number E-MTAB-1628^16^. Data were then normalized and differential expression was assessed using the Bioconductor package EdgeR. Genes with at least two-fold changes with a p value <0.05 were considered as differentially expressed. Expression data from 21,722 genes was obtained and compared between oil and E2 treatment, using two complementary approaches, including analysis of individual genes for differential expression, and GSEA for detecting coordinated changes in sets of genes representing pathways. Significance of gene expression changes between different conditions was assessed with an empirical Bayes statistic for differential expression (t-test), as implemented in limma package (R/Bioconductor). GSEA was performed following instructions (http://www.broadinstitute.org/gsea/index.jsp), using a weighted statistic, ranking by signal to noise ratio, 1000 gene-set permutations, and a custom gene set database including both the KEGG pathways (http://www.genome.jp/kegg/pathway.html) and gene list manually compiled from the literature, including the gene list comparing *Igf1r*-null HSCs with WT HSCs^16^.

### Statistics

Student’s t-test was used to calculate the *P* values unless otherwise indicated. Pearson’s correlation coefficient analysis was used to calculate the *P* values of competitive BM transplantation assay. Data were reported as mean ± SEM.

## Acknowledgements

We thank the Brigham & Women’s Hospital (BWH) Flow Cytometry Core and Yiling Qiu for assistance with FACS sorting, Dr. Stuart Orkin [Boston Children’s Hospital (BCH)] for mouse lines, and Dr. Huafeng Xie (BCH) for help with FACS analysis and irradiation. This research was supported by NIH grant R01 HL107663 (to Z.L.), AACR-Aflac, Incorporated Career Development Award for Pediatric Cancer Research (10-20-10-LI, to Z.L.), and BWH Startup Fund (to Z.L.).

## Author contributions

Y.X. designed and conducted experiments, analyzed data, and co-wrote the manuscript. D.X., X.H., H.P., E.P., J.C., D.E.L., and L.T. conducted experiments for mouse models. Z.L. designed experiments, analyzed data, and wrote the manuscript.

## References

1 Leeman, D. S. & Brunet, A. Stem cells: Sex specificity in the blood. Nature 505, 488–490 (2014). 10.1038/505488a

2 Hudry, B., Khadayate, S. & Miguel-Aliaga, I. The sexual identity of adult intestinal stem cells controls organ size and plasticity. Nature 530, 344–348 (2016). 10.1038/nature16953

3 Nakada, D. et al. Oestrogen increases haematopoietic stem-cell self-renewal in females and during pregnancy. Nature 505, 555–558 (2014). 10.1038/nature12932

4 Orkin, S. H. & Zon, L. I. Hematopoiesis: an evolving paradigm for stem cell biology. Cell 132, 631–644 (2008). S0092-8674(08)00125-6 [pii] 10.1016/j.cell.2008.01.025

5 Morrison, S. J. & Scadden, D. T. The bone marrow niche for haematopoietic stem cells. Nature 505, 327–334 (2014). 10.1038/nature12984

6 Bowie, M. B. et al. Hematopoietic stem cells proliferate until after birth and show a reversible phase-specific engraftment defect. J Clin Invest 116, 2808–2816 (2006). 10.1172/JCI28310

7 Nygren, J. M., Bryder, D. & Jacobsen, S. E. Prolonged cell cycle transit is a defining and developmentally conserved hemopoietic stem cell property. Journal of immunology 177, 201–208 (2006).

8 Pietras, E. M., Warr, M. R. & Passegue, E. Cell cycle regulation in hematopoietic stem cells. J Cell Biol 195, 709–720 (2011). 10.1083/jcb.201102131

9 Pollak, M. Insulin and insulin-like growth factor signalling in neoplasia. Nat Rev Cancer 8, 915–928 (2008).

10 Wahner Hendrickson, A. E., et al. Expression of insulin receptor isoform A and insulin-like growth factor-1 receptor in human acute myelogenous leukemia: effect of the dual-receptor inhibitor BMS-536924 in vitro. Cancer Res 69, 7635–7643 (2009). 0008-5472.CAN-09-0511 [pii] 10.1158/0008-5472.CAN-09-0511

11 Kolb, E. A. & Gorlick, R. Development of IGF-IR Inhibitors in Pediatric Sarcomas. Curr Oncol Rep 11, 307–313 (2009).

12 Zumkeller, W. The insulin-like growth factor system in hematopoietic cells. Leuk Lymphoma 43, 487–491 (2002).

13 Zhang, C. C. & Lodish, H. F. Insulin-like growth factor 2 expressed in a novel fetal liver cell population is a growth factor for hematopoietic stem cells. Blood 103, 2513–2521 (2004). 10.1182/blood-2003-08-2955 2003-08-2955 [pii]

14 Chou, S. & Lodish, H. F. Fetal liver hepatic progenitors are supportive stromal cells for hematopoietic stem cells. Proc Natl Acad Sci U S A 107, 7799–7804 (2010). 1003586107 [pii] 10.1073/pnas.1003586107

15 Garrett, R. W. & Emerson, S. G. The role of parathyroid hormone and insulin-like growth factors in hematopoietic niches: physiology and pharmacology. Mol Cell Endocrinol 288, 6–10 (2008). S0303-7207(08)00110-X [pii] 10.1016/j.mce.2008.02.022

16 Venkatraman, A. et al. Maternal imprinting at the H19-Igf2 locus maintains adult haematopoietic stem cell quiescence. Nature 500, 345–349 (2013). 10.1038/nature12303

17 Klusmann, J. H. et al. Developmental stage-specific interplay of GATA1 and IGF signaling in fetal megakaryopoiesis and leukemogenesis. Genes Dev 24, 1659–1672 (2010). 24/15/1659 [pii] 10.1101/gad.1903410

18 Dietrich, P., Dragatsis, I., Xuan, S., Zeitlin, S. & Efstratiadis, A. Conditional mutagenesis in mice with heat shock promoter-driven cre transgenes. Mamm Genome 11, 196–205 (2000).

19 Stadtfeld, M. & Graf, T. Assessing the role of hematopoietic plasticity for endothelial and hepatocyte development by non-invasive lineage tracing. Development 132, 203–213 (2005). 10.1242/dev.01558

20 Oguro, H., Ding, L. & Morrison, S. J. SLAM family markers resolve functionally distinct subpopulations of hematopoietic stem cells and multipotent progenitors. Cell Stem Cell 13, 102–116 (2013). 10.1016/j.stem.2013.05.014

21 Kuhn, R., Schwenk, F., Aguet, M. & Rajewsky, K. Inducible gene targeting in mice. Science 269, 1427–1429 (1995).

22 Fagan, D. H. & Yee, D. Crosstalk between IGF1R and estrogen receptor signaling in breast cancer. J Mammary Gland Biol Neoplasia 13, 423–429 (2008).

23 Knudsen, K. J. et al. ERG promotes the maintenance of hematopoietic stem cells by restricting their differentiation. Genes Dev 29, 1915–1929 (2015). 10.1101/gad.268409.115

24 Xie, Y. et al. Reduced Erg Dosage Impairs Survival of Hematopoietic Stem and Progenitor Cells. Stem Cells 35, 1773–1785 (2017). 10.1002/stem.2627

25 Vlaeminck-Guillem, V. et al. Mutual repression of transcriptional activation between the ETS-related factor ERG and estrogen receptor. Oncogene 22, 8072–8084 (2003). 10.1038/sj.onc.1207094

26 Sanchez-Aguilera, A. et al. Estrogen signaling selectively induces apoptosis of hematopoietic progenitors and myeloid neoplasms without harming steady-state hematopoiesis. Cell Stem Cell 15, 791–804 (2014). 10.1016/j.stem.2014.11.002

27 Oguro, H. et al. 27-Hydroxycholesterol induces hematopoietic stem cell mobilization and extramedullary hematopoiesis during pregnancy. J Clin Invest 127, 3392–3401 (2017). 10.1172/JCI94027

28 Notta, F., Doulatov, S. & Dick, J. E. Engraftment of human hematopoietic stem cells is more efficient in female NOD/SCID/IL-2Rgc-null recipients. Blood 115, 3704–3707 (2010). 10.1182/blood-2009-10-249326

29 Suman, S., Johnson, M. D., Fornace, A. J., Jr. & Datta, K. Exposure to ionizing radiation causes long-term increase in serum estradiol and activation of PI3K-Akt signaling pathway in mouse mammary gland. Int J Radiat Oncol Biol Phys 84, 500–507 (2012). 10.1016/j.ijrobp.2011.12.033

30 Clark, R. The somatogenic hormones and insulin-like growth factor-1: stimulators of lymphopoiesis and immune function. Endocr Rev 18, 157–179 (1997). 10.1210/edrv.18.2.0296

31 Medina, K. L. et al. Identification of very early lymphoid precursors in bone marrow and their regulation by estrogen. Nat Immunol 2, 718–724 (2001). 10.1038/90659

32 Medina, K. L., Strasser, A. & Kincade, P. W. Estrogen influences the differentiation, proliferation, and survival of early B-lineage precursors. Blood 95, 2059–2067 (2000).

33 Kaludjerovic, J. & Ward, W. E. The Interplay between Estrogen and Fetal Adrenal Cortex. Journal of nutrition and metabolism 2012, 837901 (2012). 10.1155/2012/837901

34 Calvanese, V., Lee, L. K. & Mikkola, H. K. Sex hormone drives blood stem cell reproduction. EMBO J 33, 534–535 (2014). 10.1002/embj.201487976

35 Li, Z., Chen, M. J., Stacy, T. & Speck, N. A. Runx1 function in hematopoiesis is required in cells that express Tek. Blood 107, 106–110 (2006).

36 Ober, C., Loisel, D. A. & Gilad, Y. Sex-specific genetic architecture of human disease. Nat Rev Genet 9, 911–922 (2008). 10.1038/nrg2415

37 Subramanian, A. et al. Gene set enrichment analysis: a knowledge-based approach for interpreting genome-wide expression profiles. Proc Natl Acad Sci U S A 102, 15545–15550 (2005).

38 Gao, J. et al. Hedgehog signaling is dispensable for adult hematopoietic stem cell function. Cell Stem Cell 4, 548–558 (2009). 10.1016/j.stem.2009.03.015

39 Hofmann, I. et al. Hedgehog signaling is dispensable for adult murine hematopoietic stem cell function and hematopoiesis. Cell Stem Cell 4, 559–567 (2009). 10.1016/j.stem.2009.03.016

40 Koch, U. et al. Simultaneous loss of beta-and gamma-catenin does not perturb hematopoiesis or lymphopoiesis. Blood 111, 160–164 (2008). 10.1182/blood-2007-07-099754

41 Maillard, I. et al. Canonical notch signaling is dispensable for the maintenance of adult hematopoietic stem cells. Cell Stem Cell 2, 356–366 (2008). 10.1016/j.stem.2008.02.011

42 Cobas, M. et al. Beta-catenin is dispensable for hematopoiesis and lymphopoiesis. J Exp Med 199, 221–229 (2004). 10.1084/jem.20031615

43 Kabiri, Z. et al. Wnts are dispensable for differentiation and self-renewal of adult murine hematopoietic stem cells. Blood 126, 1086–1094 (2015). 10.1182/blood-2014-09-598540

44 Zhao, C. et al. Loss of beta-catenin impairs the renewal of normal and CML stem cells in vivo. Cancer Cell 12, 528–541 (2007). 10.1016/j.ccr.2007.11.003

45 Zhao, C. et al. Hedgehog signalling is essential for maintenance of cancer stem cells in myeloid leukaemia. Nature 458, 776–779 (2009). 10.1038/nature07737

46 Chiang, M. Y., Radojcic, V. & Maillard, I. Oncogenic Notch signaling in T-cell and B-cell lymphoproliferative disorders. Curr Opin Hematol 23, 362–370 (2016). 10.1097/MOH.0000000000000254

47 Hawsawi, Y., El-Gendy, R., Twelves, C., Speirs, V. & Beattie, J. Insulin-like growth factor - oestradiol crosstalk and mammary gland tumourigenesis. Biochim Biophys Acta 1836, 345–353 (2013). 10.1016/j.bbcan.2013.10.005

48 Li, Z. et al. Developmental stage-selective effect of somatically mutated leukemogenic transcription factor GATA1. Nat Genet 37, 613–619 (2005).

49 Li, Z. et al. ETV6-NTRK3 fusion oncogene initiates breast cancer from committed mammary progenitors via activation of AP1 complex. Cancer Cell 12, 542–558 (2007).

