## Supplementary materials for "Interplay of IGF1R and estrogen signaling regulates hematopoietic stem and progenitor cells"

| Genotype | WBCs (x10^3 cells/ml) | RBCs (x10^6 cells/ml) | Platelet counts (x10^3/ml) |
|---|---|---|---|
| *VavCre* | 8.28±1.37 | 8.28±0.19 | 1081±22 |
| *VavCre;Igf1r*$^{L/L}$ | 6.16±0.91 | 7.92±0.69 | 955±130 |
| statistic | N=4, p=0.25 | N=4, p=0.64 | N=4, p=0.38 |

WBCs, white blood cells; RBCs, red blood cells.

Supplementary Fig. 1

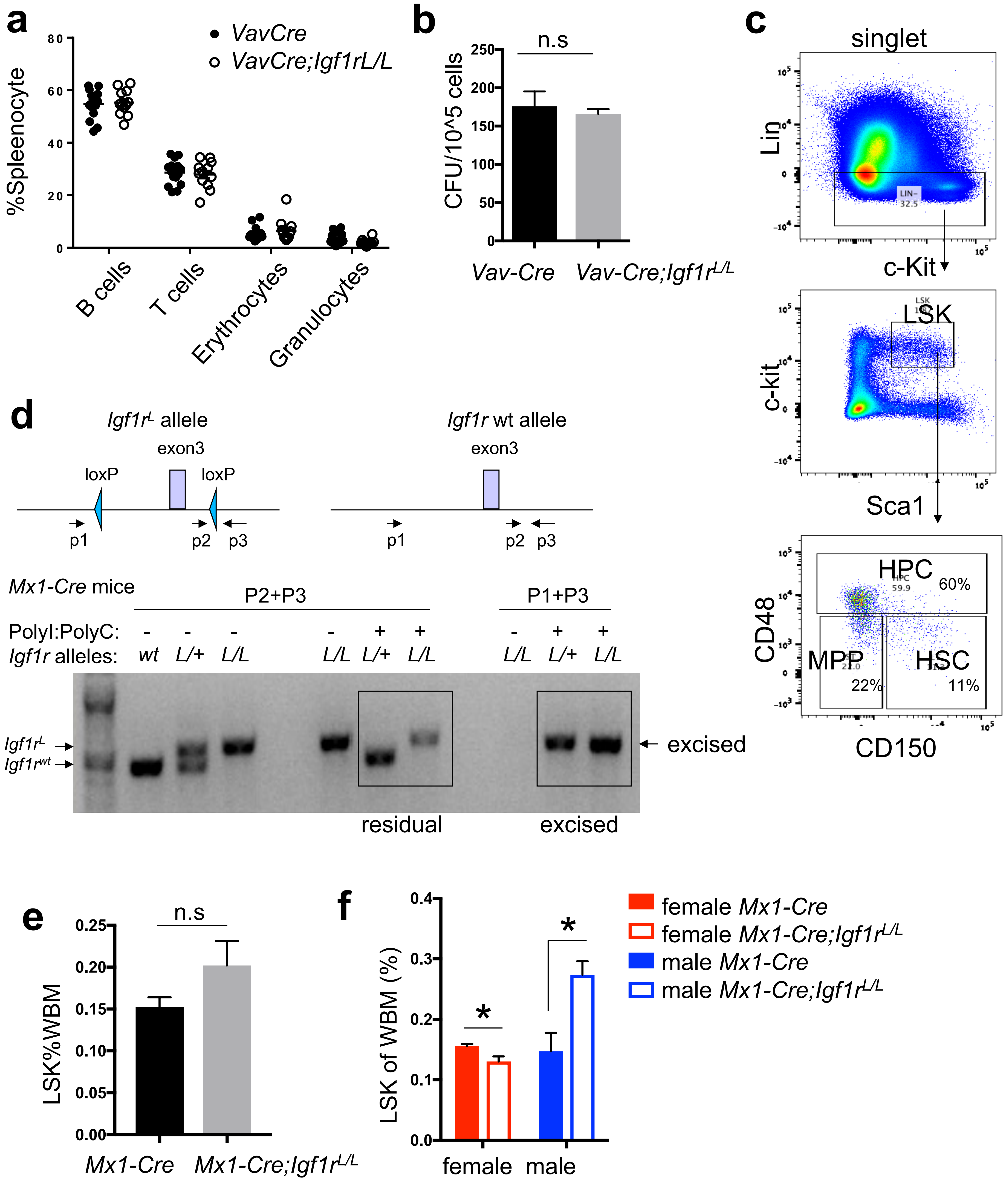

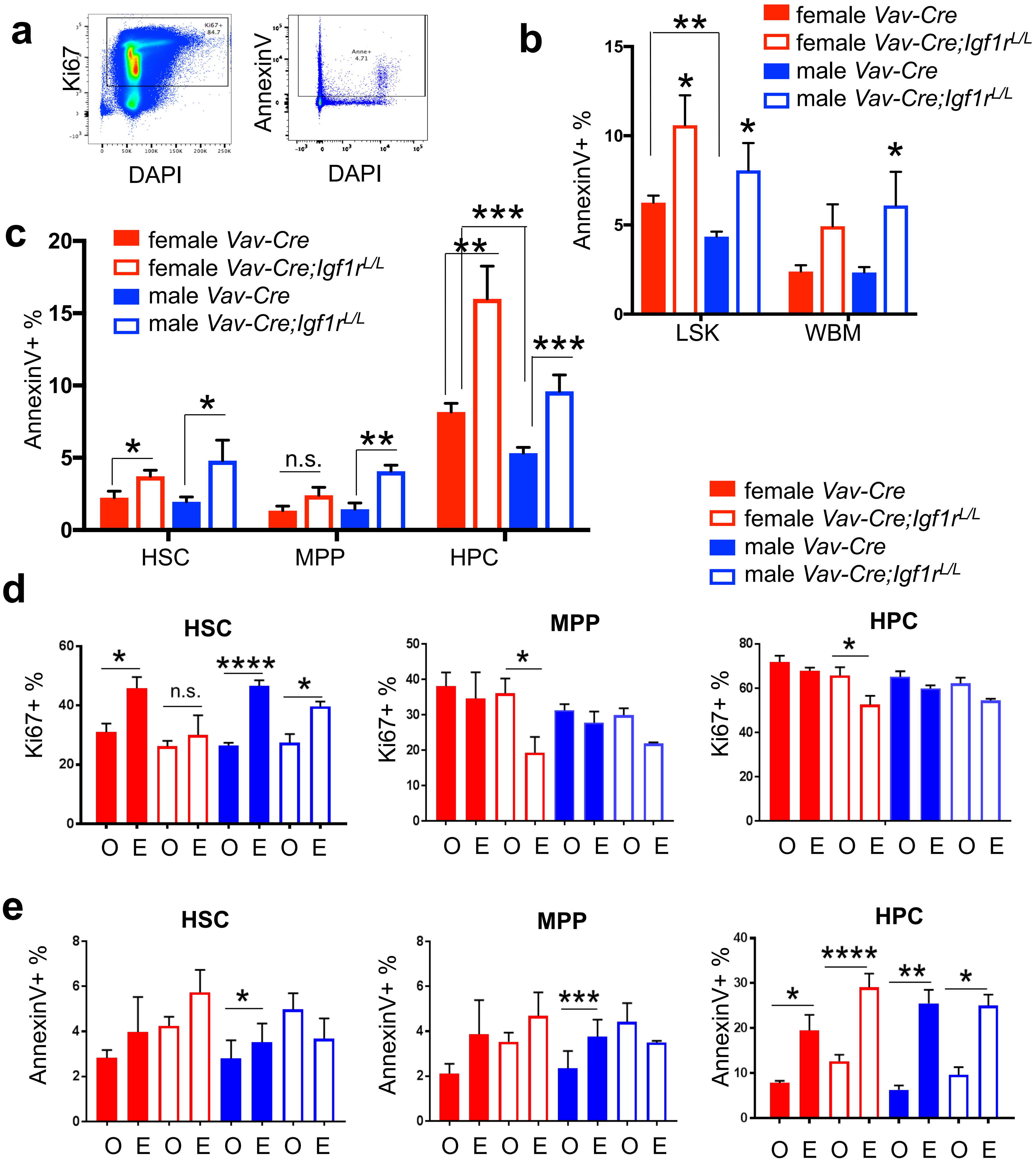
a
Ki67
DAPI
Ki67+
84.7
AnnexinV
DAPI
Anne+
4.71
b
AnnexinV+ %
LSK
WBM
**
*
*
*
female Vav-Cre
female Vav-Cre;Igf1r L/L
male Vav-Cre
male Vav-Cre;Igf1r L/L
c
AnnexinV+ %
HSC
MPP
HPC
n.s.
***
**
d
HSC
MPP
HPC
Ki67+ %
O E O E O E O E
****
e
AnnexinV+ %

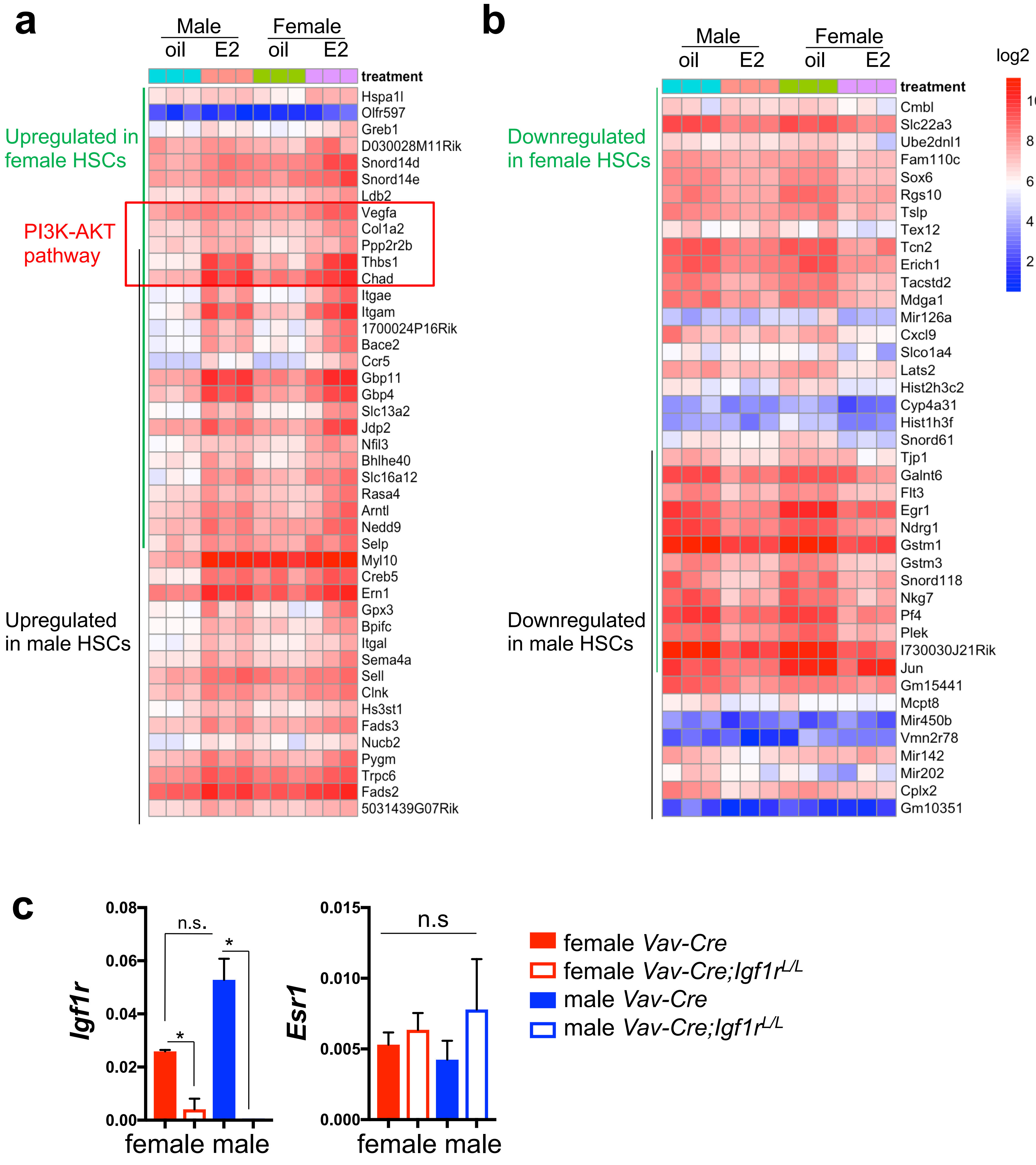
a
Male
Female
oil
E2
oil
E2
treatment
Upregulated in female HSCs
PI3K-AKT pathway
Upregulated in male HSCs
Hspa1l
Olfr597
Greb1
D030028M11Rik
Snord14d
Snord14e
Ldb2
Vegfa
Col1a2
Ppp2r2b
Thbs1
Chad
Itgae
Itgam
1700024P16Rik
Bace2
Ccr5
Gbp11
Gbp4
Slc13a2
Jdp2
Nfil3
Bhlhe40
Slc16a12
Rasa4
Arntl
Nedd9
Selp
Myl10
Creb5
Ern1
Gpx3
Bpifc
Itgal
Sema4a
Sell
Clnk
Hs3st1
Fads3
Nucb2
Pygm
Trpc6
Fads2
5031439G07Rik
b
Male
Female
oil
E2
oil
E2
treatment
log2
10
8
6
4
2
Downregulated in female HSCs
Downregulated in male HSCs
Cmbl
Slc22a3
Ube2dnl1
Fam110c
Sox6
Rgs10
Tslp
Tex12
Tcn2
Erich1
Tacstd2
Mdga1
Mir126a
Cxcl9
Slco1a4
Lats2
Hist2h3c2
Cyp4a31
Hist1h3f
Snord61
Tjp1
Galnt6
Flt3
Egr1
Ndrg1
Gstm1
Gstm3
Snord118
Nkg7
Pf4
Plek
I730030J21Rik
Jun
Gm15441
Mcpt8
Mir450b
Vmn2r78
Mir142
Mir202
Cplx2
Gm10351
c
Igf1r
0.08
0.06
0.04
0.02
0.00
n.s.
*
*
female male
Esr1
0.015
0.010
0.005
0.000
n.s
female male
female Vav-Cre
female Vav-Cre;Igf1r L/L
male Vav-Cre
male Vav-Cre;Igf1r L/L

Supplementary Fig. 4

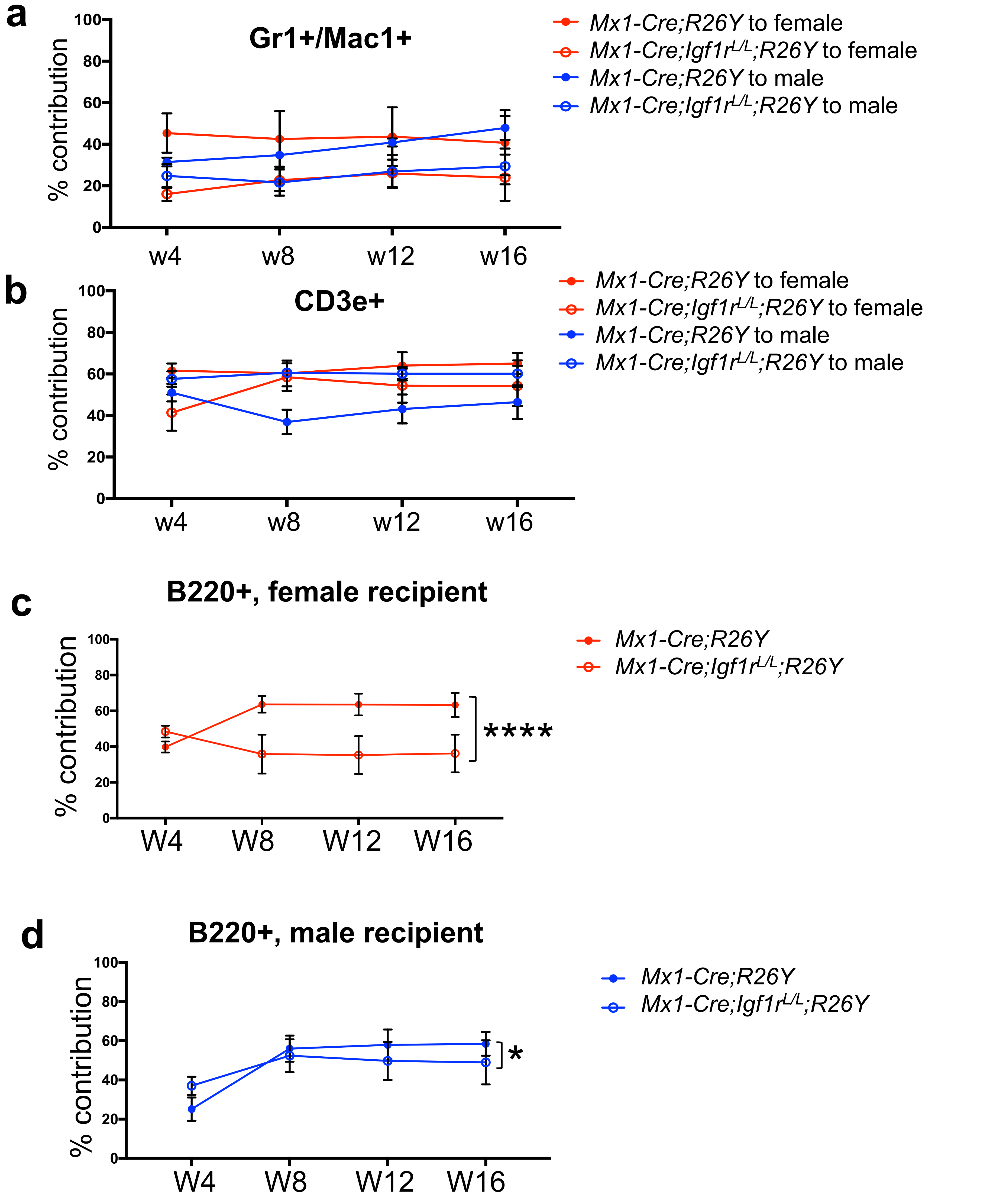

**a**

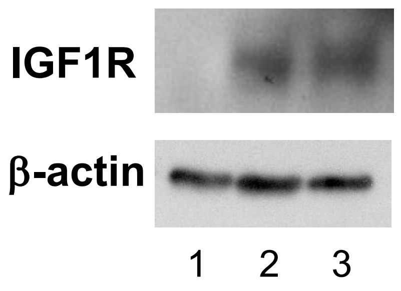


1. *Tie2-Cre;Igf1r$^{L/L}$*
2. *Tie2-Cre;Igf1r$^{L/+}$*
3. *Tie2-Cre*

**b**

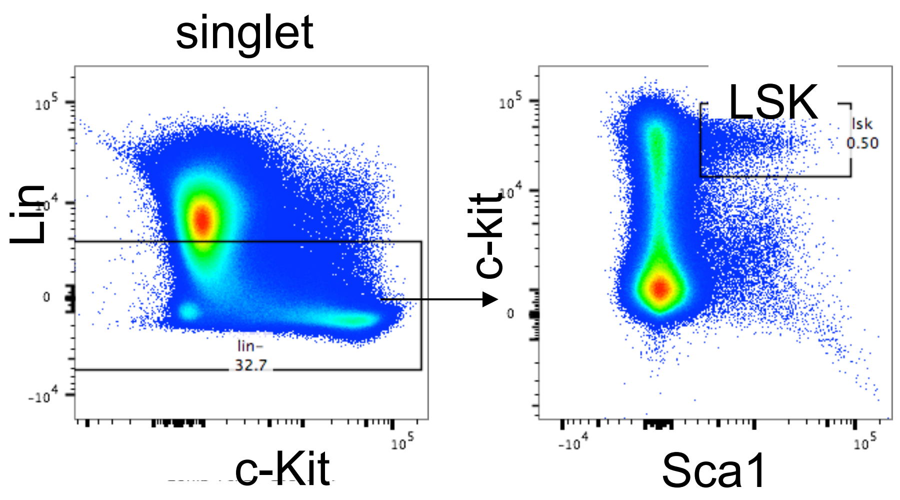


**c**

| **Genotype** | ***Tie2-Cre; Igf1r$^{L/L}$*** | ***Tie2-Cre; Igf1r$^{L/+}$*** | ***Tie2-Cre; Igf1r$^{+/+}$*** |
|---|---|---|---|
| Live pups observed (%) | 38 (22%) | 91 (52%) | 47 (27%) |
| Expected % | 25% | 50% | 25% |

Supplementary Fig. 6

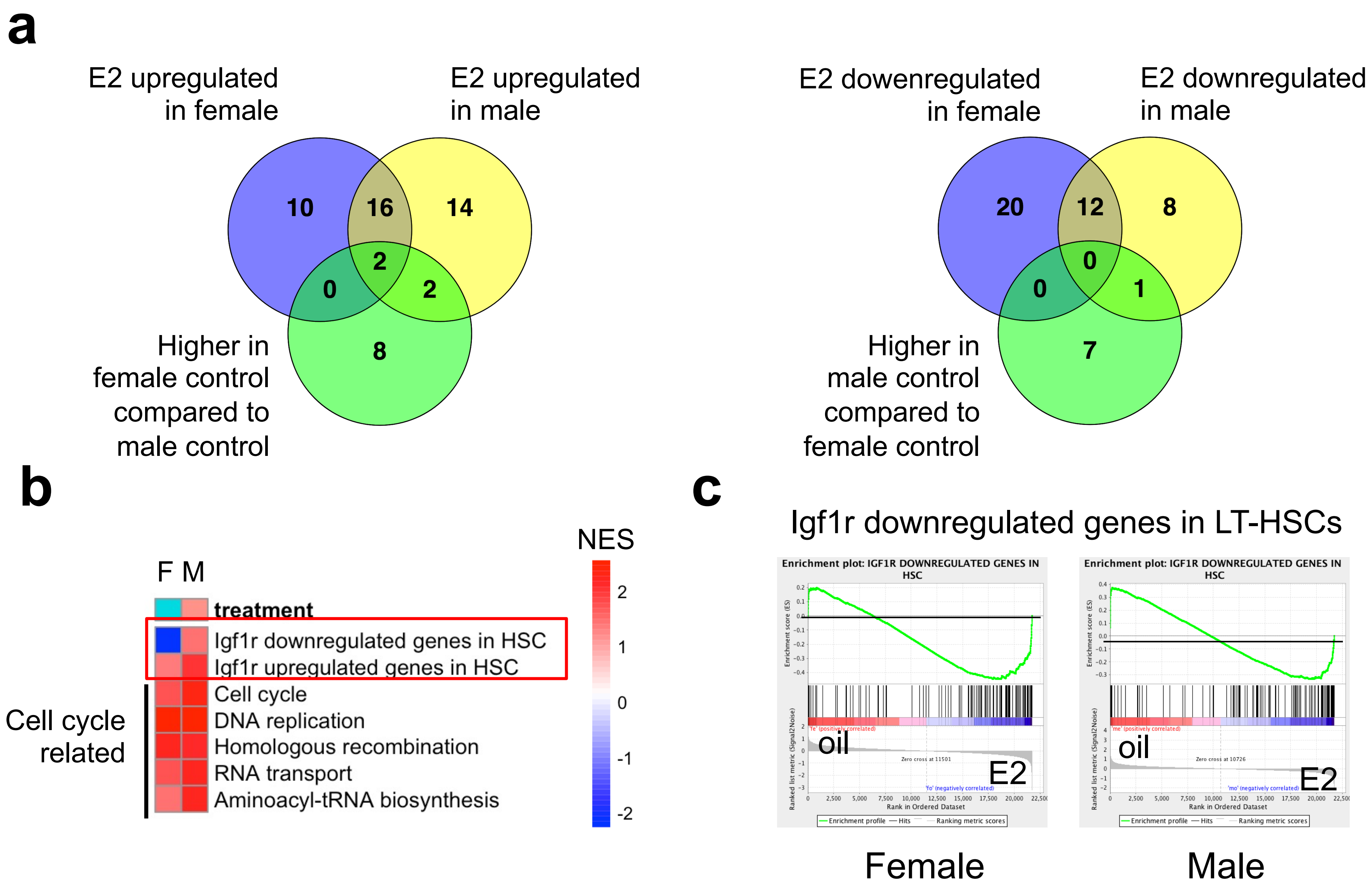

### Supplementary Figure Legends

**Supplementary Fig. 1. IGF1R signaling is largely dispensable for adult hematopoiesis.**

(a) Percentages of different lineages of mature hematopoietic cells in the spleen (based on spleenocytes) were measured by flow cytometry ($B220^+$ B cells, $CD3e^+$ T cells, $Ter119^+$ erythrocytes, $Gr1^+/Mac1^+$ granulocytes) between wild-type (WT) (*Vav-Cre*, n=14) and mutant (*Vav-Cre;Igf1r*$^{L/L}$*,* n=12). (b) Colony forming units (CFU) of BM cells were measured. n=5 for both *Vav-Cre;Igf1r*$^{L/L}$ and *Vav-Cre* (WT) mice. (c) Representative FACS plots showing strategy to determine percentage of each population. (d) Schematic diagram and PCR results showing PCR strategy for detecting *loxP* sites floxing the *Igf1r* exon3 and Cre-mediated excision of this exon. (e) The LSK population in whole bone marrow (WBM) exhibited no significant difference between *Mx1-Cre* (WT) and *Mx1-Cre;Igf1r*$^{L/L}$ (n=7 and n=8, respectively) adult mice. (f) LSK population in the BM of female or male *Mx1-Cre;Igf1r*$^{L/L}$ mice compared to *Mx1-Cre* control mice. n=4 for both female *Mx1-Cre;Igf1r*$^{L/L}$ and *Mx1-Cre* mice, n=4 and n=3 for both male *Mx1-Cre;Igf1r*$^{L/L}$ and *Mx1-Cre* mice, respectively. Data represent mean ± SEM. *$p<0.05$, **$p<0.01$, ***$p<0.005$, ****$p<0.001$, n.s.=not significant.

**Supplementary Fig. 2. Both IGF1R-loss and estrogen treatment induce apoptosis in HSPCs.**

(a) Representative FACS plots showing strategy to determine percentages of $Ki67^+$ cycling cells and $AnnexinV^+$ apoptotic cells. (b) Percentages of $AnnexinV^+$ apoptotic cells were measured for both LSK and WBM populations. $n\geq3$. (c) Percentages of $AnnexinV^+$ apoptotic cells were measured for all HSPC subpopulations. $n\geq3$. (d-e) Percentages of $Ki67^+$ proliferating cells (d) and $AnnexinV^+$ apoptotic cells (e) in each indicated HSPC subpopulation from female or male *Vav-*

*Cre;Igf1r*$^{L/L}$ and/or *Vav-Cre* mice treated with oil (O) or β-estradiol (E). n≥3. Data represent mean ± SEM. *p<0.05, **p<0.01, ***p<0.005, ****p<0.001, n.s.=not significant.

**Supplementary Fig. 3. Molecular analysis of HSCs.**

(a-b) Genes significantly upregulated (logFC>1, p<0.05, a) or downregulated (logFC<-1, p<0.05, b) by E2-treatment compared to oil-treatment in female or male HSCs respectively. Green line: genes significantly changed in female HSCs; black line: genes significantly changed in male HSCs. Red and blue indicate gene expression levels (from highest to lowest) from original microarray data. (c) qRT-PCR analysis showing relative expression levels of *Igf1r* and *Esr1* in female and male BM HSCs with the indicated genotypes, both normalized to *Gapdh*. n≥3. Data represent mean ± SEM. *p<0.05, **p<0.01, ***p<0.005, ****p<0.001, n.s.=not significant.

**Supplementary Fig. 4. Engraftment of the same donor cells to female or male recipients.**

(a and b) Donor contribution (as percentage of CD45.2$^{+}$YFP$^{+}$ cells) over time in Gr1$^{+}$/Mac1$^{+}$ granulocyte (a) and CD3e$^{+}$ T cells (b) lineages was measured in both female and male recipient mice. (c-d) Donor contribution (as percentage of CD45.2$^{+}$YFP$^{+}$ cells) in peripheral blood in the B220$^{+}$ B cell lineage was measured over time in female (c) versus male (d) recipient mice. n≥3 for each data point. Data represent mean ± SEM. Pearson's correlation coefficient analysis was used to calculate the *P* values for the whole trend. *p<0.05, ****p<0.001, n.s.=not significant.

**Supplementary Fig. 5. Analysis of IGF1R-loss in fetal hematopoiesis.**

(a) Western blot showing loss of IGF1R protein in the fetal liver (FL) hematopietic cell- derived colonies from the *Tie2-Cre* model. (b) Representative FACS plots showing strategy to determine percentages of LSK population in the FL. (c) The expected Mendelian ratio and observed ratio of mice with the indicated genotype after birth.

**Supplementary Fig. 6. Additional molecular analysis of HSCs.**

(a) Venn diagram for E2 co-regulated genes as well as genes differentially expressed in male and female HSCs (all comparison logFC>1 or <-1, p<0.05). Left panel: co-upregulated genes and genes higher in female HSCs; Right panel: co-downregulated genes and genes higher in male HSCs. (b) Representative gene sets significantly regulated (NES>1 or <-1, FDR<0.15) in both male and female HSCs after E2 treatment. Red and blue indicate gene sets significantly up- or down-regulated, respectively, in HSCs from E2-treated mice. The color scale represents their normalized enrichment score (NES). (c) Gene set enrichment analysis of Igf1r negatively correlated genes in LT-HSCs in the E2 regulated genes in female (left panel) or male (right panel) HSCs.
